# Attention differentially affects acoustic and phonetic feature encoding in a multispeaker environment

**DOI:** 10.1101/2020.06.08.141234

**Authors:** Emily S. Teoh, Edmund C. Lalor

## Abstract

Humans have the remarkable ability to selectively focus on a single talker in the midst of other competing talkers. The neural mechanisms that underlie this phenomenon remain incompletely understood. In particular, there has been longstanding debate over whether attention operates at an early or late stage in the speech processing hierarchy. One way to better understand this is to examine how attention might differentially affect neurophysiological indices of hierarchical acoustic and linguistic speech representations. In this study, we do this by using encoding models to identify neural correlates of speech processing at various levels of representation. Specifically, using EEG recorded during a “cocktail party” attention experiment, we show that phonetic feature processing is evident for attended, but not unattended speech. Furthermore, we show that attention specifically enhances isolated indices of phonetic feature processing, but that such attention effects are not apparent for isolated measures of acoustic processing. These results provide new insights into the effects of attention on different pre-lexical representations of speech, insights that complement recent anatomical accounts of the hierarchical encoding of attended speech. Furthermore, our findings support the notion that – for attended speech – phonetic features are processed as a distinct stage, separate from the processing of the speech acoustics.

## Introduction

The ability to focus on a single talker amidst multiple sounds is essential for human communication. Cherry (1953) brought this phenomenon to the fore, investigating our behavioral capacity for selective auditory attention. His study stimulated subsequent inquiries and the proposal of various theories of attention (Broadbent, 1958; Moray, 1959; Deutsch & Deutsch, 1963; Treisman 1964; Johnston & Wilson, 1980). These theories model attention as a selective filter that rejects the unattended message beyond a certain information processing stage. However, whether this filter operates at an early or late stage of speech processing is still unresolved.

In recent years, neuroscience has sought to contribute to this debate by leveraging our increased understanding of the hierarchical nature of speech processing in the human cortex. In particular, to construe meaning from sound, the cortex is posited to compute several intermediate levels of increasingly abstract representations in different functionally specialized regions of a hierarchically organized cortical network (Hickok & Poeppel, 2007; Peelle et al., 2010; Rauschecker & Scott, 2009; DeWitt & Rauschecker, 2012; Kell et al., 2019). For example, single talker neuroimaging studies have revealed that core auditory regions code low-level features which are combined in higher areas to yield more abstract neural codes (Binder et al., 2000; Davis & Johnsrude, 2003; Okada et al., 2010). And in terms of multi-talker selective attention, studies have shown that primary auditory cortical responses represent all talkers irrespective of attentional state (O’Sullivan et al., 2019), but in “higher” areas such as the superior temporal gyrus (STG), only the attended speaker is represented (Mesgarani and Chang, 2012; Zion Golumbic et al., 2013; O’Sullivan et al., 2019). Additionally, EEG/MEG studies examining the latency of neural responses (which can be considered as a rough proxy measure of processing at different hierarchical stages) have shown that all talkers are co-represented in early components, with distinct responses to the attended speaker appearing only in later components (Ding & Simon, 2012; Power et al., 2012; Puvvada & Simon, 2017). Importantly, this specificity is not something that is necessarily true for more simplistic non-speech stimuli and tasks (e.g., Power et al., 2011).

As well as understanding how attention affects processing in different brain areas and at different latencies, we also wish to understand how attention influences the encoding of the different speech *representations* that are thought to be computed by this hierarchical network. In recent years, encoding/decoding methods (Crosse et al., 2016; Holdgraf et al., 2017) have revealed neural indices of a number of these representations – from low-level acoustics to semantics – in brain responses to continuous natural speech (Ahissar et al., 2001; Brodbeck et al., 2018; Broderick et al., 2018; Daube et al., 2019; Di Liberto et al., 2018; Di Liberto et al., 2015; Mesgarani et al., 2014; Tang et al., 2017; Teoh et al., 2019). And it has been shown that neural indices of lexical (Brodbeck et al., 2018) and semantic (Broderick et al., 2018) processing can only be found for attended speech, in contrast to acoustically-driven measures like those based on the amplitude envelope that are less affected by attention (Brodbeck et al., 2018). Taken together, the results from these anatomical, time-resolved, and representational approaches support the notion that higher-order regions (and higher-level representations) are more greatly modulated by attention.

In this study, we explore how attention modulates neural indices of speech processing at *prelexical* levels. Specifically, we show that EEG reflects phonetic feature processing of attended – but not unattended – speech. We also show clear attention effects on isolated measures of phonetic feature processing, but find no such effects on isolated measures of acoustic processing. This builds on the idea of hierarchical attention effects suggested by previous studies by showing a sharp dissociation in the influence of attention on different aspects of sublexical processing. Furthermore, it supports the notion of an acoustic-phonetic prelexical processing stage in cortex, something that has been proposed in recent years (Di Liberto et al., 2015; Khalighinejad, et al., 2017; Brodbeck et al., 2018; Gwilliams et al., 2020; Mesgarani et al., 2014), but that has also been challenged (Daube et al., 2019).

## Methods

### Subjects

Fourteen subjects (nine female and five male) between the ages of 19 and 30 participated in the experiment. All subjects were right-handed and spoke English as their primary language. Subjects reported no history of hearing impairment or neurological disorder. Each subject provided written informed consent prior to testing and received monetary reimbursement. The study was approved by the Research Subjects Review Board at the University of Rochester. Some of the data used here (the first 10 trials) were analyzed differently for a previous study (Teoh & Lalor, 2019).

### Stimuli and Procedures

Subjects undertook 40 one-minute trials in two separate blocks. Stimuli consisted of two works of fiction, one narrated by a female talker and the other by a male talker. Silent gaps in the audio exceeding 0.3 s were truncated to 0.3 s in duration.

The stimuli were filtered using Head-Related Transfer Functions (HRTFs), giving rise to the perception that the talkers were at 90 degrees to the left and right of the subject. The HRTFs used were obtained from the CIPIC database (Algazi et al., 2001). Subjects were always instructed to attend to one of the two talkers – a counterbalanced paradigm was employed in which, over the course of the experiment, they would have attended to both male and female talkers and at both locations.

Before the experiment, subjects were asked to minimize motor activities and to maintain visual fixation on a crosshair centered on the screen during trials. After each trial, subjects were required to answer four multiple-choice comprehension questions on each of the stories (attended and unattended).

All stimuli were sampled with a frequency of 44 100 Hz and were presented using Sennheiser HD650 headphones and Presentation software from Neurobehavioral Systems (http://www.neurobs.com).

### Data Acquisition and Pre-processing

The experiment was conducted in a soundproof room. A Biosemi ActiveTwo system was used to record EEG data from 128 electrode positions on the scalp as well as two electrodes over the mastoid processes (all digitized at 512Hz).

EEG data were re-referenced to the mastoids. Automatic bad channel rejection and interpolation was performed. A particular channel was deemed as bad if the standard deviation of the channel was lower than a third of or exceeded three times the mean of the standard deviation of all channels. In place of the bad channel, data were interpolated from the four nearest neighboring electrodes using spherical spline interpolation (Delorme & Makeig, 2004). To decrease subsequent processing time, data were downsampled to 128Hz. Data were filtered between 0.2 – 8 Hz using a Chebyshev II filter. Low delta-band frequencies (down to 0.2 Hz) were included as they have previously been found to also be important for speech processing (Teoh et al., 2019).

### Speech Representations

We computed acoustic and phonetic speech representations of both the attended and unattended stories (Figure 1A depicts these representations for an excerpt taken from one of the audio clips).

**Figure 1:**
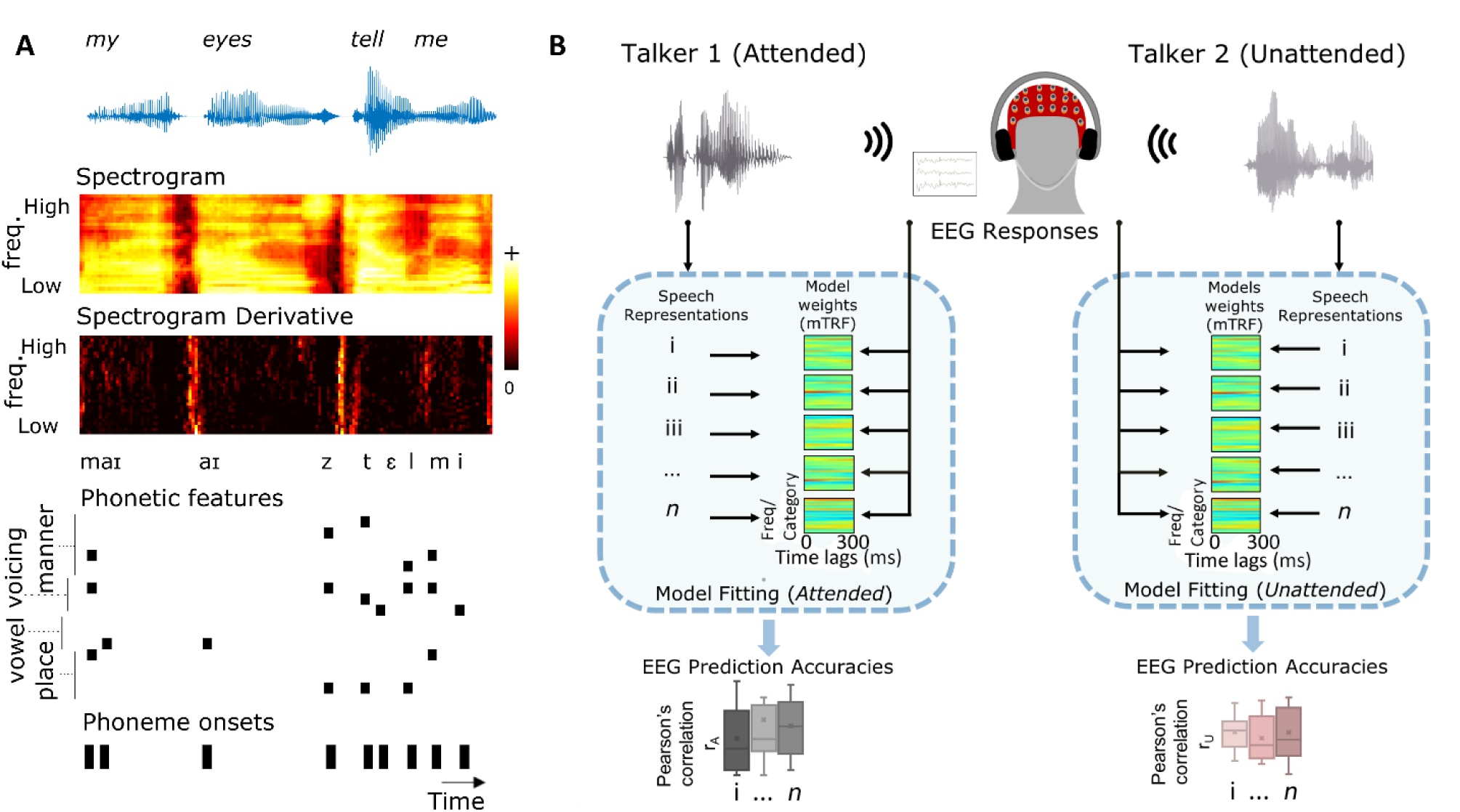
(A) Speech representations: The first row depicts an acoustic waveform of an excerpt taken from one of the stimuli. Subsequent rows show the computed acoustic and phonetic representations for that excerpt. (B) Analysis framework: Cross-validation is used to train forward models mapping the different attended and unattended speech representations to EEG. These models are then used to predict left-out EEG. Pearson’s correlation is used to evaluate model accuracy, and the prediction accuracies of acoustic and phonetic models are compared.

#### Acoustic

- Spectrogram (*s*): The GBFB toolbox (Schädler & Kollmeier, 2015) was used to extract the spectral decomposition of the time-varying stimulus energy in 31 mel-spaced bands with logarithmic compressive nonlinearity. The spectrogram representation computed in this way is consistent with Daube et al. (2019), where it was shown to better predict neural activity than other techniques.
- Spectrogram Derivative (*sD*): The temporal derivative of each spectrogram channel was computed, and then half-wave rectified. This represents an approximation of the onsets in the acoustics and was shown to contribute to predicting neural responses in previous research (Daube et al., 2019).

#### Phonetic

- Phonetic features (*f*): Phoneme-level segmentation of the stimuli was first computed using FAVE-Extract (Rosenfelder et al, 2014) and the Montreal Forced Aligner (McAuliffe et al, 2017) via the DARLA web interface (Reddy & Stanford, 2015). The phoneme representation was then mapped into a space of 19 features (based on the University of Iowa’s phonetics project). The features describe the articulatory and acoustic properties of the phonetic content of speech. Based on the onset time of each feature, a multivariate time-series binary matrix (19 features by samples) was produced.
- Phonetic feature onsets (*fo*): Univariate vector of the onsets of all phonetic features (i.e., a summary measure of all onsets in *f* without categorical discrimination). This was included as an additional control. Specifically, if the *f* representation above fails to improve EEG prediction beyond the *fo* representation, it would suggest that the encoding of individual phonetic features is not reflected in EEG data.

### Model fitting: Multivariate Linear Regression

The acoustic and phonetic representations for both attended and unattended speech described above were normalized and mapped to the concurrently recorded 128-channel EEG signals. Multivariate regularized linear regression was employed to relate the features to the recorded EEG data, where each EEG channel is estimated to be a linear transformation of the speech features over a range of time lags. This transformation is described by the temporal response function (TRF). For a particular speech feature, this operation can be represented mathematically as following (Crosse et al., 2016):

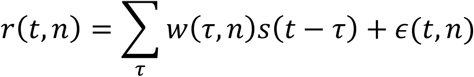

where *r*(*t, n*) is the neural (EEG) response at channel, *n* and time point, *t* = 1 … *T, s*(*t − τ*) is the multivariate stimulus representation at a lag, *τ, w*(*τ, n*) is the transformation (TRF) of the stimulus at lag *τ*, and *ϵ*(*t, n*) is the residual response not explained by the model.

The TRF is estimated by minimizing the mean square error between the actual neural response, *r*(*t, n*), and the response predicted by the transformation, 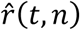. In practice, this can be solved by using reverse correlation. We use the mTRF toolbox (Crosse et al., 2016 – https://sourceforge.net/projects/aespa/), which solves for the TRF (*w*) using reverse correlation with ridge regression:

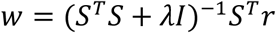

where *λ* is the ridge regression parameter, *I* is the identity matrix, and the matrix *S* is the lagged time series of the stimulus matrix, *s*. The TRF approach can be used to relate multiple features of the stimuli to the ongoing EEG simultaneously by extending the lagged stimulus matrix to include the various features (more details can be found in Crosse et al., 2016). The ridge regression parameter was tuned using leave-one-out cross validation. That is, we trained on *n*-1 trials for a wide range of *λ* values (e.g. 1, 1×10^1^, 1×10^2^, … 1×10^9^), computed the average TRF across trials for each *λ*, then tested the TRFs on the *n*th trial. This was repeated *n* times, rotating the trial to be tested each time. The *λ* value that maximized the Pearson’s correlation coefficient between the actual and predicted neural response over all trials was selected.

The transformation was computed over a range of time lags, reflecting the idea that changes in the features of the ongoing stimulus are likely to produce effects in the ongoing EEG in that interval. Specifically, we chose the interval 0 to 300 ms based on previous EEG-based speech studies, where no visible response was present outside this range when considering EEG responses to acoustic and phonetic features (Di Liberto et al., 2015; Lalor & Foxe, 2010). We quantified how well each speech representation related to the neural data using leave-one-trial-out cross-validation (as described above), with Pearson’s correlation coefficient as our metric of prediction accuracy. Because the cross-validation procedure takes the average of the validation metric across trials, the models are not biased toward the test data used for cross-validation (Crosse et al., 2016).

To evaluate whether a feature contributed independently of all other features in predicting the neural data, we also computed the partial correlation coefficients (Pearson’s *r*) between the EEG predicted by each measure’s model with the actual recorded EEG after controlling for the effects of all other features.

### Statistical Testing

To test that a partial correlation coefficient is above chance level, we performed non-parametric permutation testing. The predicted EEG activity for each model’s representation was permuted across trials such that they were matched to the actual EEG of a different trial, and partial correlation coefficients were computed, controlling for the effects of all other features. This was done 1000 times for each subject to establish a distribution of chance-level prediction accuracies. To perform group-level statistical testing, we generated a null distribution of group means: one prediction accuracy from each subject’s individual distribution was selected at random to go into each group mean. This process was repeated 1000 times, sampling with replacement for each subject. For comparison between groups (e.g. attended vs unattended for a particular representation), two-tailed Wilcoxon signed-rank testing was used.

## Results

Subjects were found to be compliant in carrying out the behavioral task (Figure 2A). The average questionnaire accuracy was 73.2 ± 3.2% when subjects were tested on the attended story and 27.2 ± 1.6% for unattended stimuli (theoretical chance level is 25% as there are four possible answers to each question).

**Figure 2.**
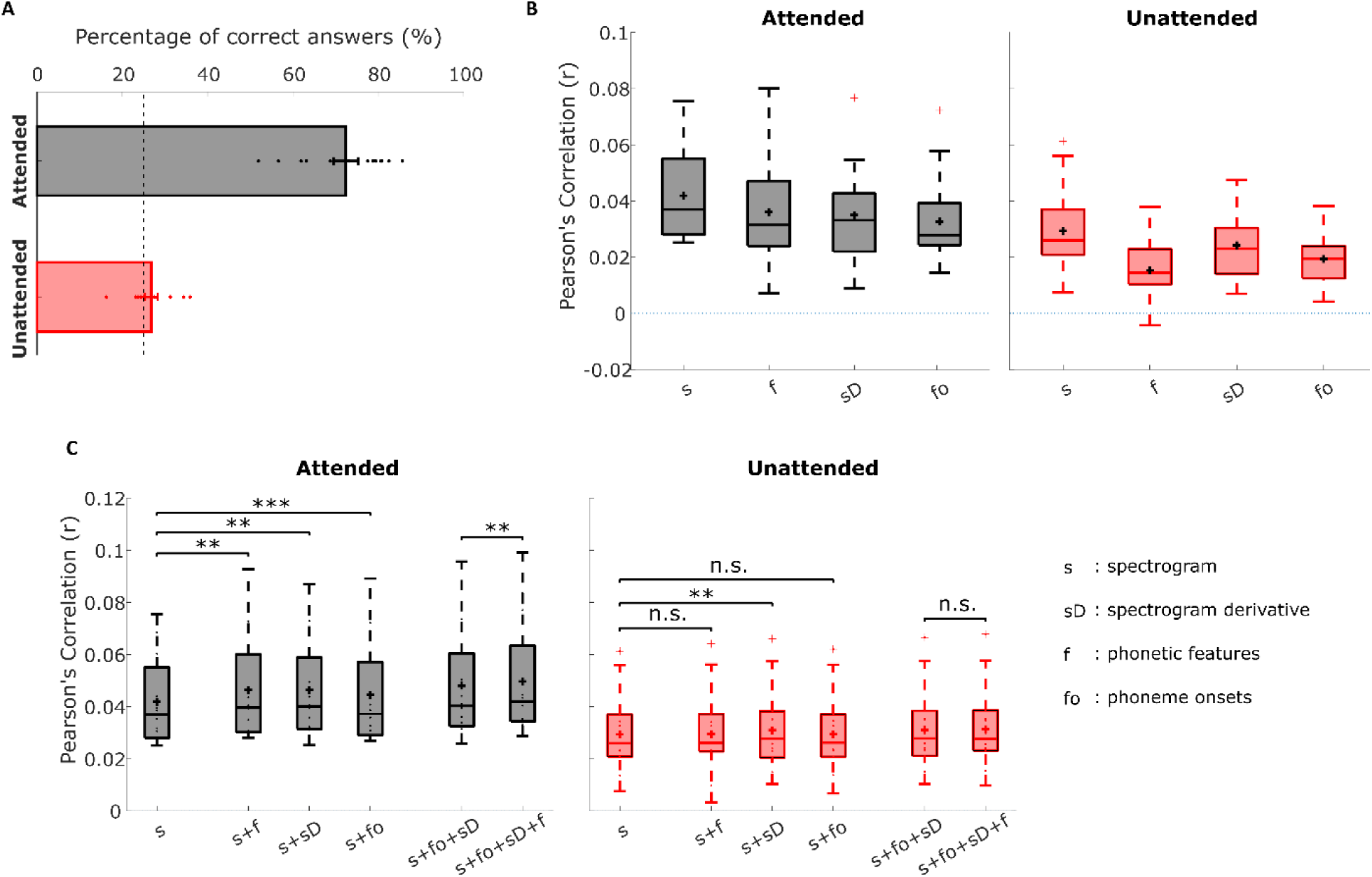
(A) Behavioral (comprehension questionnaire) results – dots indicate individual subject performance. Theoretical chance level is 25% (multiple-choice test with four options) and is indicated by the dashed line (B) Prediction accuracies of individual acoustic and phonetic feature spaces (as labelled on horizontal axes) for the attended and unattended stimuli. There is redundancy between these features, so joint modelling was performed to gauge in particular whether phonetic features adds unique predictive power beyond other features (C) Prediction accuracies of joint feature spaces for attended and unattended stimuli (**p<0.01, two-tailed Wilcoxon sign-ranked test).

### Responses to the attended talker reflect phonetic-level representations

We assessed the performance of attended and unattended models that were trained to predict EEG responses using the acoustic and phonetic feature representations extracted from the speech stimulus (Figure 2B). The Pearson’s correlation between predicted and actual EEG activity, averaged across the same 12 bilateral fronto-temporal channels used in Di Liberto et al. (2015) and Daube et al. (2019), was used as our metric of prediction accuracy for each model (although the same pattern of results was observed if all 128 channels were used). All individual features could predict EEG above chance on a group level (non-parametric permutation test, p=0.999e-3).

Now, the feature spaces used here are not independent, so the individual model predictions are not a pure representation of the extent to which EEG tracks these features. Phonetic features (*f*) overlaps with the spectrogram (*s*) in that if every phoneme is always spoken the same way, then the two representations would be equivalent. The phoneme onset (*fo*) representation marks the same time-points as the phonetic feature representation (*f*), except that it does not contain feature category information. The spectrogram derivative (*sD*), an approximation of acoustic onsets, contains peaks that overlap with those of the phoneme onset (*fo*) representation. Thus, the processing of a particular feature was assessed by examining whether adding that feature improved prediction of the EEG responses beyond a model built from other features (Figure 2C). We were particularly interested in establishing whether the three other features could improve prediction accuracy beyond using spectrograms alone, and if phonetic features contributed unique predictive power beyond all other features. To this end, we performed pairwise statistical testing of the joint model prediction accuracies – these results are shown in Table 1.

**Table 1.** Joint model comparison statistics: Wilcoxon two-sided signed rank test results for attended and unattended stimuli (columns). Each set of rows test a different statistical question, as specified by the baseline. Terms being evaluated are to the left of the ‘>’ symbol. Bolded values indicate significant improvement over baseline

|  | Attended | Unattended |
| --- | --- | --- |
| baseline: $s$ | | |
| $s + f > \text{baseline}$ | **$p = 0.0015, z = 3.1702$** | $p = 0.3003, z = 1.0358$ |
| $s + sD > \text{baseline}$ | **$p = 0.0023, z = 3.0447$** | **$p = 0.0019, z = 3.1074$** |
| $s + fo > \text{baseline}$ | **$p = 9.82\text{e-}04, z = 3.2958$** | $p = 0.1578, z = 1.4125$ |
| baseline: $s + sD$ | | |
| $s + sD + f > \text{baseline}$ | **$p = 0.0043, z = 2.8563$** | $p = 0.1981, z = 1.2869$ |
| baseline: $s + fo$ | | |
| $s + fo + f > \text{baseline}$ | **$p = 0.0023, z = 3.0447$** | $p = 0.6378, z = 0.4708$ |
| baseline: $s + fo + sD$ | | |
| $s + fo + sD + f > \text{baseline}$ | **$p = 0.0076, z = 2.6680$** | $p = 0.4703, z = 0.7219$ |

We found that including phonetic features significantly improved prediction over the acoustic feature spaces when subjects were attending to the stimuli; but this improvement was not observed for the unattended stimuli. Importantly, it also improved prediction over phoneme onsets in the attended case, indicating that the inclusion of specific phonetic feature categories carries unique and important information. The acoustic onset (i.e., spectrogram derivative) representation displayed a different pattern of results – including this measure along with the spectrogram (i.e., *s+sD*) improved prediction accuracy over spectrogram alone for both attended and unattended stimuli, suggesting that it is less modulated by attention.

### Attention enhances isolated measures of phonetic feature processing, but not acoustic processing

Given the aforementioned redundancies between the predictions of feature spaces, we were also interested in more clearly isolating the unique contribution of each feature. To do so, we employed a partial correlation approach to control for the predictions of all other representations. The unique predictive power of each model is shown in Figure 3A (shown here for an average across 12 channels, although including all 128 channels revealed the same pattern of results). On a group level, all features made unique, significant contributions to the EEG predictions except the spectrogram derivative models (attended and unattended), and the unattended phonetic features model (non-parametric permutation test; sD attended: p=0.0619; sD unattended: p=0.1788; f unattended: p=0.1788; all other models: p<0.05). When comparing attended and unattended models, we found a significant effect of attention for phonetic features (two-tailed Wilcoxon signed rank; p<0.05), but not for any of the other representations. The topographic distributions for the unique predictive power of each model are shown in Figure 3B.

**Figure 3.**
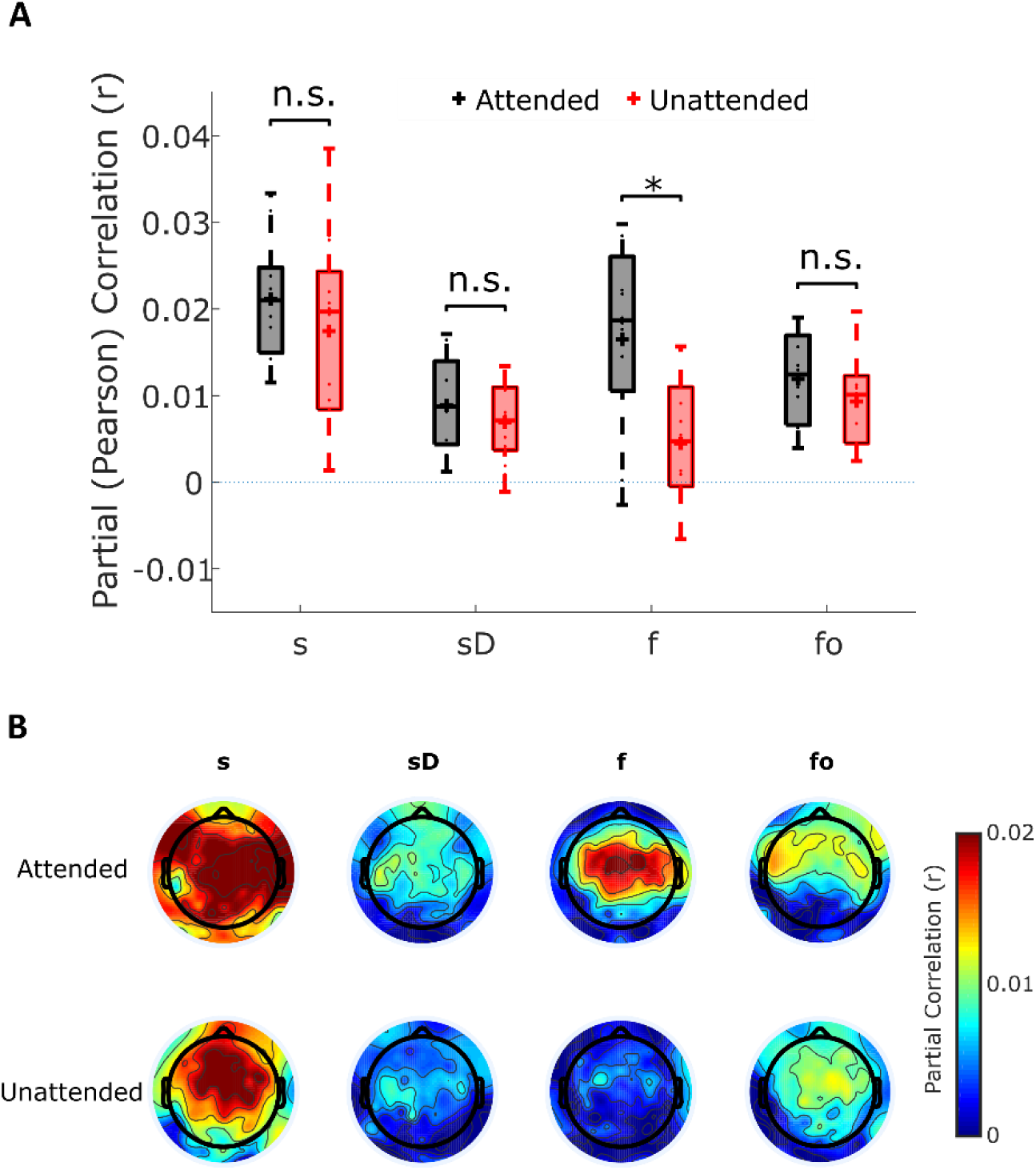
(A) Unique predictive power for the acoustic and phonetic feature spaces under two attentional states. The features are as labelled on the horizontal axis. Statistical testing was carried out to identify attentional effects (two-tailed Wilcoxon signed rank; *p<0.05). (B) Topographic distribution of partial correlations, averaged across all subjects.

### Phonetic feature categories contribute predictive power beyond differentiating vowels and consonants

Di Liberto et al. (2015) found a high degree of discriminability between EEG responses to vowels and consonants. The phonetic feature representation used in the above analysis information on the manner, place, and voicing of consonants, as well as the articulatory position of vowels. We wondered how our results would be affected if – instead of using a 19-dimensional vector – the phonemes were simply marked as vowels or consonants. We repeated the partial correlation analysis described above and found that there was a significant decrease in prediction accuracy when the information on specific articulatory features was left out in the case of attended speech, but not for unattended speech (two-tailed Wilcoxon signed-rank test, p=0.011, z=2.542). Additionally, there was no longer a significant difference between the prediction accuracies of attended and unattended stimuli (att_vc vs. unatt_vc; Wilcoxon signed-rank test; p = 0.4326; z = 0.7847)

## Discussion

There has been longstanding debate on how selective attention affects the processing of speech. Here, we set out to investigate attentional modulation at the pre-lexical level. In particular, based on earlier work (Di Liberto et al. 2015; Daube et al., 2019), we considered how two acoustic and two phonetic feature representations of attended and unattended speech were reflected in EEG responses to that speech. We found that, for attended speech, including a 19-dimensional phonetic features representation improved the prediction of the EEG responses beyond that obtained when only using acoustic features. This was not true for unattended speech. Furthermore, we found that the unique predictive power of the phonetic features representation was enhanced for attended versus unattended speech. This was not true for any other feature. We contend that these findings make two important contributions to the literature. First, they contribute to the debate around pre-lexical speech processing in cortex. And second, they contribute to the longstanding debate on how selective attention affects the processing of speech

In terms of prelexical speech processing, our study shows that a phonetic feature representation has unique predictive power when it comes to modelling responses to attended speech (Fig 2 & Table 1). This suggests that processing attended, but not unattended speech, may involve a mapping from an acoustic to a categorical phonemic representation. This is a controversial idea. In particular, while linguistic theories and prominent speech processing models posit a mapping to phonetic features and/or phonemes as an intermediate stage between low-level acoustics and words (McClelland & Elman, 1986; Liberman et al., 1967; Hickok & Poeppel, 2007), this viewpoint is not unanimous. Some researchers question the necessity and existence of a mental representation at this level for comprehending speech meaning (Lotto & Holt, 2000). Indeed, there have been theories advocating for alternatives, including a different intermediate representational unit (e.g. syllables; Massaro, 1974), or direct matching to the lexicon based on the comparison of acoustic representations (Goldinger, 1998; Klatt, 1989). In support of an acoustic-phonemic stage of processing, there has been evidence from behavioral and EEG studies that humans perceive phoneme categories. Liberman et al. (1957) found that discrimination within phoneme categorical boundaries is poorer than across them. Eimas et al. (1971) looked at infants’ ability to discriminate syllables that differed in a voicing (phonetic) feature and found that infants also perceived categorical phonetic features. Additionally, Näätänen et al. (2001) found that the mismatch negativity (MMN), an event-related potential component that reflects discriminable change in some repetitive aspect of the ongoing auditory stimulation, is observed when the deviant stimulus is a phoneme prototype of a subject’s native language relative to when it is a non-prototype. However, it has been argued that these studies used simple isolated units, and therefore their results could be due to the type of task and not reflective of everyday listening.

More recent neuroimaging studies have shown that a phonetic feature representation of speech can predict neural activity during listening to natural, continuous speech. Using high-density cortical surface electrodes, Mesgarani et al. (2014) found that the superior temporal gyrus (STG) selectively responds to phonetic features (their Figure 2-4). Di Liberto et al. (2015) and Khalighinejad et al. (2017) found evidence that non-invasive EEG activity reflects the categorization of speech into phonetic features. However, a recent paper challenged these findings and argued that neural responses to speech could be well explained by a model that is based entirely on acoustic features (Daube et al., 2019). Namely, they found that acoustic onsets made very similar predictions to the benchmark phonetic features, and that acoustic onsets explained parts of the neural response that phonetic features could not. Here, we show evidence to support the encoding of a rich array of phonetic features (certainly more than simply vowel and consonant categorization; Fig. 4) for attended speech. It is possible that we were able to find unique predictive power for these phonetic features because our study involved stimuli from two different talkers (a male and a female) while the stimuli used in Daube et al. (2019) were spoken by a single talker. As mentioned earlier, there is redundancy between speech acoustics and a phonetic feature representation because each phonetic feature has a characteristic spectro-temporal acoustic profile. This overlap can make it difficult to isolate the precise contribution of each predictor. Indeed, if there were no variation in spectro-temporal acoustics across each utterance of the same phoneme, the phonetic feature and spectrogram representations would be, effectively, identical. However, the more variation there is between utterances of the same phoneme (which will happen when including both male and female speakers), the greater the difference should be between the contribution of neurons that care about those variations (e.g., low-level “acoustic” neurons) and the contribution of neurons whose responses are invariant to the acoustic details and only dependent on phoneme category. As such, it may be that, when using stimuli with high intra-phoneme acoustic variation, including both acoustic and phonetic feature representations is of most benefit for predicting neural activity. Future work will investigate this idea further.

**Figure 4.**
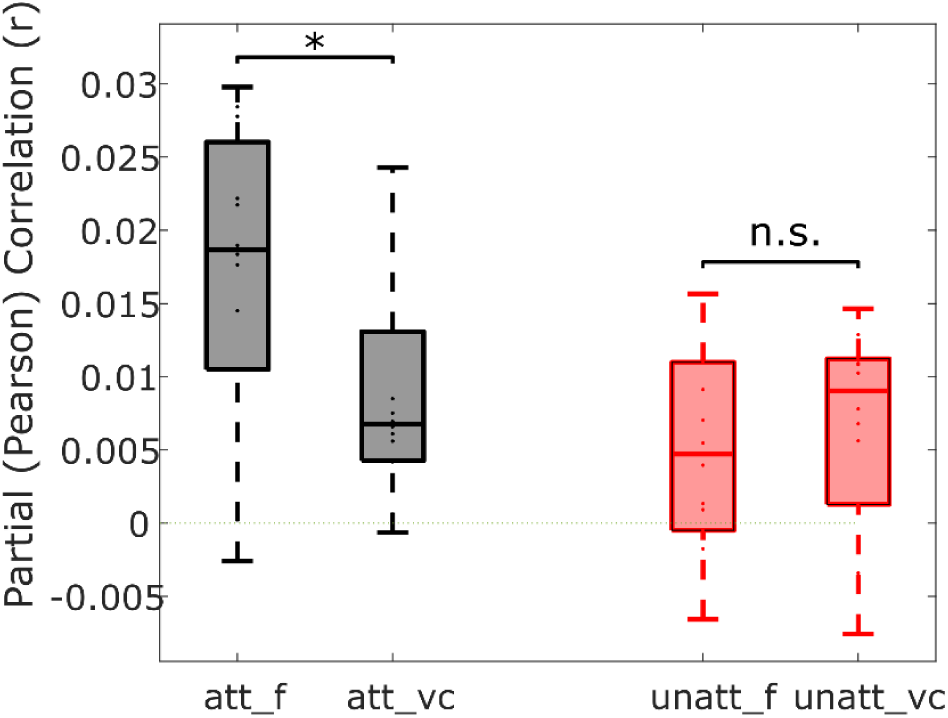
A significant reduction in unique predictive power was observed for the attended condition when the partial correlation analysis was repeated with phonemes only categorised as vowels or consonants (vc), as opposed to their underlying articulatory features (f) (two-tailed Wilcoxon signed-rank test, *p=0.011, z=2.542).

Of course, there are also other acoustic transformations of the auditory stimulus that have not been considered beyond the spectrogram and its derivative, as well as other theorized intermediate representations apart from phonemes/phonetic features that have not been tested. It is likely that some of these feature sets may overlap with our measure of phonetic features. As such, we cannot definitively prove the encoding of this measure in cortex. However, one reason the idea of mapping to words via an acoustic-phonetic stage has prevailed is due to its computational efficiency (Scharenborg, Norris, Bosch, & McQueen, 2005). It is less obvious how the detection of acoustic onsets and other acoustic representations would lead to lexical access without an intermediate stage – and in particular, how such a model would robustly generalize to new words and new speakers.

As to the cocktail party attention debate, our results suggest that attention differentially modulates cortical processing of acoustic and phonetic information. We found that the *unique* predictive power of phonetic features was modulated by attention, while the other feature spaces were not. This result is in line with a wealth of evidence for stronger attention effects at higher levels of the speech processing hierarchy. For example, there is no doubt that both attended and unattended speech will influence activity in the cochlea and the auditory nerve. But by the time one reaches auditory association areas like STG, only the attended speaker is represented (Mesgarani and Chang, 2012; Zion Golumbic et al., 2013). Indeed, an important recent study used simultaneous recordings in both primary auditory cortex (Heschl’s gyrus; HG) and STG, to show that both attended and unattended speech are robustly represented in primary areas, with STG selectively representing the attended speaker. This lines up very well with our results. In particular, it is well established that spectro-temporal models of sounds can do a good job of explaining neural responses in primary auditory areas, but not in nonprimary auditory cortex (Norman-Haignere & McDermott, 2018). Thus, we might expect our isolated acoustic speech processing indices to mostly reflect activity from primary auditory areas, which have been shown to be relatively unaffected by attention (O’Sullivan et al., 2019). Meanwhile STG is known to be selective for phonetic feature representations (Mesgarani & Chang, 2014) and activity in STG is strongly affected by attention (O’Sullivan et al., 2019). Thus, we might expect our isolated measures of phonetic feature processing to be strongly influenced by activity from STG, and thus, significantly affected by attention. This is the pattern of results that we have observed.

Additional support for our findings as evidence for a higher-level locus of attentional selection comes from research examining the latency of attention effects on EEG and MEG responses to speech. This approach has commonly focused on a rather general measure of speech processing that derives from indexing how the neural data track the speech envelope. Directly determining how much this speech processing is driven by acoustic versus speech-specific processing is not clear, although there is undoubtedly a very substantial acoustic contribution (Lalor et al., 2009; Howard and Poeppel, 2010; Prinsloo & Lalor, 2020). Nonetheless, researchers have examined components of these responses at different latencies as a proxy measure of processing at different hierarchical stages, and found that both attended and unattended speech are well represented in early components, with distinct responses to the attended speaker appearing only in late components (Ding & Simon, 2012; Power et al., 2012; Puvvada & Simon, 2017). Directly linking envelope tracking at different latencies with different representations of speech is a task for future research. Indeed, it may be quite an important task given the myriad efforts in recent years aimed at linking the effects of attention on behavior with those on a difficult-to-interpret neurophysiological measure that is likely substantially driven by attentionally-insensitive acoustic processing (e.g., Tune et al., 2020). Similarly, it may be that decoding of attentional selection from neural recordings will be improved by focusing on how those recordings reflect specific speech representations that are strongly affected by attention rather than the very general speech envelope (O’Sullivan et al., 2015).

Finally, it is important to add the caveat that our lack of an attention effect on isolated measures of acoustic processing is not evidence of their absence. It may well be that our analysis pipeline is not sensitive enough to detect small attention effects on those measures. Nonetheless, the fact that a clear difference was found for the phonetic feature representation supports the notion of stronger attention effects at higher levels of the speech processing hierarchy. The capacity of the brain to process information from multiple streams at any given moment in time is limited, so a longstanding question has concerned the extent to which unattended speech undergoes processing in cortex. Our results, considered together with studies focused on lexical (Brodbeck et al., 2018) and semantic (Broderick, et al., 2018) processing, suggest it possible that attention is categorically selective for speaker-invariant representations and, at most, attenuates lower-level acoustic (speaker-dependent) measures.

